# Galaxy-SynBioCAD: Synthetic Biology Design Automation tools in Galaxy workflows

**DOI:** 10.1101/2020.06.14.145730

**Authors:** Melchior du Lac, Thomas Duigou, Joan Hérisson, Pablo Carbonell, Neil Swainston, Valentin Zulkower, Forum Shah, Léon Faure, Mostafa Mahdy, Paul Soudier, Jean-Loup Faulon

**Affiliations:** Micalis Institute, INRAE, AgroParisTech, University Paris-Saclay, Jouy-en-Josas, France; Genomique Metabolique, Genoscope, Institut Francois Jacob, CEA, CNRS, Univ Evry, University Paris-Saclay, 91057, Evry, France; Manchester Institute of Biotechnology, SYNBIOCHEM center, School of Chemistry, The University of Manchester, Manchester M1 7DN, UK; Instituto Universitario de Automatica e Informatica Industrial, Universitat Politecnica de Valencia, 46022 Valencia, Spain; Institute of Systems, Molecular and Integrative Biology University of Liverpool, Liverpool L69 7ZB, UK; Edinburgh Genome Foundry, SynthSys, School of Biological Sciences, University of Edinburgh, EH93BF Edinburgh, UK

**Keywords:** Design Automation, Biosynthetic Pathway Engineering, Galaxy workflows, Standards, Web Application

## Abstract

Many computer-aided design tools are available for synthetic biology and metabolic engineering. Yet, these tools can be difficult to apprehend, sometimes requiring a level of expertise that limits their use by a wider community. Furthermore, some of the tools, although complementary, rely on different input and output formats and cannot communicate with one another. Scientific workflows address these shortcomings while offering a novel design strategy. Among the workflow systems available, Galaxy is a web-based platform for performing findable and accessible data analyses for all scientists regardless of their informatics expertise, along with interoperable and reproducible computations regardless of the particular platform that is being used.

Here, we introduce the Galaxy-SynBioCAD^a^ portal, the first Galaxy toolshed for synthetic biology and metabolic engineering. It allows one to easily create workflows or use those already developed by the community. The portal is a growing community effort where developers can add new tools and users can evaluate the tools performing design for their specific projects. The tools and workflows currently shared on the Galaxy-SynBioCAD portal cover an end-to-end metabolic pathway design process from the selection of strain and target to the calculation of DNA parts to be assembled to build libraries of strains to be engineered to produce the target.

Standard formats are used throughout to enforce the compatibility of the tools. These include SBML for strain and pathway and SBOL for genetic layouts. The portal has been benchmarked on 81 literature pathways, overall, we find we have a 65% (and 88%) success rate in retrieving the literature pathways among the top 10 (50) pathways predicted and generated by the workflows.

## Introduction

Computation has become an essential tool in life science research. Synthetic biology is no exception to that trend. While historically synthetic biology was mostly focused on the rational design of genetic logic devices from modular DNA parts, it is now being developed and used for biotechnological widespread applications, including the design of metabolic pathways for the production of chemicals. A fundamental goal of synthetic biology is to make biological systems easier to engineer. As part of this endeavor, significant attention is being paid to the development of workflows that will assist researchers through the synthetic biology lifecycle. Disregarding the application, synthetic biology consistently follows a Design-Build-Test-Learn workflow and adheres to design principles from engineering such as standardization and abstraction of modular parts, as well as the decoupling of design from fabrication in order to speed up the process.

Following the electronic design automation (EDA) concept, which was an essential contribution that spurred the digital society we live in, there are many design automation tools for circuit and pathway, these are extensively reviewed in Appleton *et al*.^1^ and Lin *et al*.^2^ respectively. As an example, Cello^3^ applies the EDA approach to genetic circuits. With Cello, a specific desired logic function is encoded into the Verilog language (a standard hardware description language used to model and design electronic circuits) which in turn is transformed into a linear DNA sequence that can be constructed and eventually run in living cells. The user enters the desired logic function and a “user constraints file” which contains the details on a logic gate library, the layout of the genetic system, the organism and strain, and the operating conditions for which the circuit design is valid. Additionally, Cello encompasses a combinatorial algorithm allowing to design multiple constructs containing the same circuit while varying unconstrained design elements to build a library that can be screened. Cello eventually includes a simulator generating predicted cytometry distribution for all combinations of input states, which can be directly compared to flow cytometry experiments. Cello was applied to the design of 60 circuits for *Escherichia coli*, 45 (75%) of which performed correctly in every output state.

Cello comprises several steps, which are connected and therefore need to use standardized input/output formats. Among those formats are Verilog to represent a logic function, JSON to describe the user constraints, and Eugene^4^ to encode a set of parts and constraints between the parts. While Cello achieved to compile and standardize several pieces of software for genetic design, in general, available Synthetic Biology design tools are far from parallel that achievement. Yet, two main standards have emerged in the past two decades. The first, SBML^5^ is a biological modeling standard that has been developed by the systems biology community and is currently supported by more than 250 different software tools. The primary goal of SBML is to enable exchange between modeling and simulation software for biological systems like for instance metabolic pathways and networks. The second standard, SBOL^6^, is a data exchange standard specific to synthetic biology. SBOL has been developed to document genetic components (DNA, RNA, protein, etc.) for the purpose of engineering design. SBOL can now encode complex genetic circuits, metabolic pathways, vectors, and plasmids.

One of the biggest challenges and a barrier to the reuse of successful designs, is that biological data relevant to the design of novel systems are often not exchanged. Addressing this challenge, the SynBioHub repository^7^ is an open-source software project that facilitates the sharing of information about engineered biological systems using the SBOL format. SynBioHub provides computational access for software and data integration, and a web-based graphical user interface that enables users to search for and share designs.

As pointed in Appleton *et al*.^1^, most of the tools mentioned above feature dedicated support for designing genetic regulatory networks/circuits, but they do not feature the same level of support for designing biosynthetic/metabolic pathways. One of the purposes of the Galaxy-SynBioCAD portal is addressing this shortcoming by providing a suite of interoperable and standardized tools to design pathways from the design specification (choice of the compound, strain) to the DNA parts to be assembled.

As for metabolic circuit design, there are plenty of pathway design software tools^2^. Briefly, from a given target compound and a given chassis strain, the first step consists of finding metabolic reactions that are heterologous to the chassis and link the target compound to the native metabolites of the host organism. This step is carried out by retrosynthesis software^8–13^ and requires the use of reaction rules^14^ if one wishes to search for novel pathways or find pathways that produce unnatural target compounds. The result of retrosynthesis software tools is a metabolic map and there is a need in a second step to enumerate the pathways linking the chassis metabolites to the target. There are many tools for pathway enumeration and search^15^, which are sometimes integrated into the retrosynthesis software itself. The third step is to find the most promising enzyme sequences catalyzing the metabolic reactions of the enumerated pathways. This can be achieved either through similarity search to enzyme annotated metabolic reactions^16–18^, or machine learning trained on metabolic databases^19,20^. Once the pathways have been annotated with enzyme sequences, they can be ranked in a fourth step. The ranking criteria are diverse, they can be among others based on thermodynamics^21^, predicted yield of the target^22^, target rate of production through flux balance analysis^9,11,21^, chassis cytotoxicity of the target and intermediates^21^, along with simpler criteria like pathway length. Moreover, there are multiple layout solutions and settings available in order to engineer the top-ranked pathways. Indeed the individual genes coding for the enzyme can be placed under different promoters, in a different order, with different RBS strength (if the chassis is a bacteria), and on different plasmids with different origins of replication if the engineering is performed on a plasmid. The fifth step deals with this issue by making use of tools such as the RBS calculator^23^ to compute RBS sequences for different strengths, and design of experiments (DoE)^24,25^ to sample the space of possible constructs, which can be quite large. The result of that step is a library of layouts representing either the same or different pathways. At this stage one can either synthesize the whole layout DNA or, as it is most commonly done, synthesize individual DNA parts. Several computational tools can be used to perform this sixth and last step before engineering the pathways, these tools compute parts to be synthesized depending on the assembly protocol chosen by the user. With the DNA parts in hands engineering can begin. Computation tools to help the build tasks are more sparse than for design. One can cite here Aquarium^26^, which provides instructions to a person or a robot to perform the assembly tasks along with Antha^27^, BioBlocks^28^, and DNA-BOT^29^. As with SynBioHub for designs there are repositories where protocols for the build task can be stored^30^. Engineered pathways are generally evaluated using HPLC or mass spectrometry analyses. Here too, computational tools can help in particular the workflows produced by OpenMS^31^ or Worlflow4Metabolomics^32^ and data depot^33^ exist to upload the results along with commercial data management systems like Benchling or Ryffin.

Considering the above, we are clearly at a stage where the pathway engineering process is not that far from being fully driven by computer software products. However, there are several hurdles that prevent this from happening even for tools covering pathway design only. First, the tools are not easily findable, they are stored in different places and unless you are an expert, the keywords to search online are not obvious. Secondly, some of the tools are difficult to access some requiring registration, purchase or access fees. Thirdly, almost none of the tools are interoperable and cannot be chained one after another to ensure that computational experiments are communicated well, and hence reproducible. Lastly, and perhaps most problematic for wider acceptance, the tools can be difficult to comprehend requiring a level of expertise that limits their use by a large community.

Scientific workflows help to address these issues by providing an open, web-based platform for performing findable and accessible data analyses linked to experimental protocols for all scientists regardless of their informatics expertise, along with interoperable and reproducible computations regardless of the particular platform that is being used.^34^ Indeed, without programming or informatics expertise, scientists that need to use computational approaches are impeded by difficulties ranging from tool installation to determining which parameter values to use, to efficiently combining and interfacing multiple tools together in an analysis chain. Scientific workflows provide solutions where data is combined and processed into a configurable, structured set of steps that implements computational solutions to a scientific problem. Existing systems often provide graphical user interfaces to combine different technologies along with efficient methods for using them, and thus increase the efficiency of the scientists using them. Additionally, workflow systems generally provide a platform for developers seeking a wider audience and broad integration of their tools, and can thus drive forward further developments in a specific field of research. Among existing workflow platforms, Galaxy is a system originally developed for genome analysis^35^ which now includes several thousand tools that can be found in the public ToolShed^36^.

Here, we introduce the Galaxy-SynBioCAD portal, the first Galaxy set of tools for synthetic biology and metabolic engineering. It allows one to easily create workflows or use already developed shared workflows. The portal is a growing community effort where developers can add new tools and users can evaluate the tools performing design for their specific projects. The tools and workflows currently shared on the Galaxy-SynBioCAD portal cover an end-to-end metabolic pathway design process from the selection of strain and target to the calculation of DNA parts to be assembled to build libraries of strains to be engineered to produce the target.

## Results

### Tools implementation into Galaxy nodes

To be implemented into Galaxy, software applications were selected among the computational tools mentioned in the Introduction section. Several criteria were used for this selection, (i) the tools needed to be relevant for pathway design, (ii) be published, (iii) open-source under MIT, GNU GPL, or related licenses, (iv) well documented and deposited in GitHub, (v) making use of standard input/output, and (vi) amenable to compartmentalization in Docker and implementation into a Galaxy node.

The process used to integrate computational tools into Galaxy nodes is described in the Methods section (see IT Architecture subsection where an example is provided for the tool RetroPath2.0). The list of Galaxy nodes provided below are currently installed on the Galaxy-SynBioCAD portal and enable one to design pathways from target and strain selection to DNA part calculation.

**RetroRules**^b,14^ is a searchable database of reaction rules. Reaction rules are generic descriptions of (bio)chemical reactions encoded into the community standard SMARTS. The use of reaction rules allows estimating the outcomes of chemical transformation based on the generalization of reactions available in knowledge DBs such as BRENDA^37^, MetaCyC^38^, Rhea^39^, or MetaNetX^40^. The degree of generalization is controlled by describing the surrounding environment of the reaction center up to a given diameter. To ensure the accuracy of the predicted transformations that will outcome from the reaction rules, the RetroRules dataset provided by the Galaxy RetroRules node has been validated by (i) checking that rules allow to reproduce the template reactions, and by (ii) checking that results obtained by decreasing diameters are supersets of results obtained with higher diameters. Only the reaction rules that successfully passed the 2 checks are retained. The RetroRules dataset provided presently is tagged as `rr02` and is freely downloadable from the RetroRules database^c^. The validation of this dataset has a success rate of 99.3%. The node outputs a CSV file of reaction rules in SMARTS format.

**RetroPath2.0**^d^ is an open-source tool for building retrosynthesis networks by combining reaction rules and a retrosynthesis-based algorithm to link the desired target compound to a set of available precursors^10^. Typically, the target compound, also named “source compound” is the compound of interest one wishes to produce, while the precursors are usually compounds that are natively present in a chassis strain. Starting from the source compound at the first iteration, the reaction rules matching the chemical structure of the source are applied and newly predicted chemicals are generated. For each reaction a score is calculated based on the ability to retrieve enzyme sequences catalysing substrate to product transformations. Newly produced chemicals are scanned and kept for the next iteration if they are not within the set of available precursors. In that way, a new iteration is started using the previously collected chemicals as the new source set. The iterative process stops when either no new chemicals are discovered or the predefined number of steps is reached. The node takes as input three CSV files, one with a list of sink molecules using the standard InChI format, another with a single source (target) molecule in InChi format too, and a last file containing the reaction rules in SMARTS format. The retrosynthesis network is outputted as a CSV file providing reactions in the reaction SMILES format and chemicals in both SMILES and InChI formats along with other information like the score for each reaction.

**RP2paths**^e^ in an open-source tool dedicated to the enumeration of heterologous pathways that lie in a retrosynthesis network as produced by RetroPath2.0^10^. Such analysis is a required step in our workflow to ensure that only pathways fulfilling all the precursor needs are retained for further analysis. Quickly, the main steps performed are (i) the scope reduction, which aims to reduce the size of the input metabolic network using an iterative node removal approach to retain only reactions and chemicals involved in at least one producible pathway, (ii) the stoichiometric matrix build of the subnetwork containing only the scope, *i.e*. the chemicals and reactions retained at the previous step, followed by (iii) the Elementary Flux Mode Enumeration (EFM)^41^, from which only the enumerated modes linking the target compound to precursors are output as a heterologous pathway. The node takes as input a retrosynthesis network in the CSV file produced by RetroPath2.0, and outputs the enumerated pathways (using IDs) as well as structure of involved chemicals (as SMILES) in CSV files as well.

#### Pathways to SBML and Complete Reactions

The node *Pathways to SBML*^f^ converts the output of RP2paths, as well as each individual pathway, to distinct SBML files. Those output pathways are “enriched” with additional information (see Method section) that cannot be easily stored as part of a normal SBML file and include structural information for chemical species (SMILES, InChI and InChIKey) and for each reaction a rule ID, a score based on enzyme availability produced by RetroPath2.0, and the rule itself in SMARTS format. The tools also adhere to the MIRIAM annotation standard for the cross-references of chemical species to public databases^42^. The tool takes the CSV outputs of RP2paths as well as the output of RetroPath2.0 and outputs a collection of SBML files compressed in a TAR file. The second node, called *Complete Reactions*^g^, adds the required cofactors to complete the reactions. Indeed, due to the nature of RetroPath2.0 retrosynthesis algorithm, the reactions it produces are mono-component^10^. To complete reactions, the node queries the MetaNetX database for the appropriate cofactors and adds them to the SBML files. The node takes for input either a single SBML file, or a collection compressed in a TAR. The node produces a collection of SBML files compressed in a TAR file.

**Thermodynamics**^h^ calculates the Gibbs free energy of reactions and heterologous pathways by considering every chemical species involved in each reaction. This is done using the tool eQuilibrator^43^ calculating the formation energy either using public database ID reference (when recognized with the tools internal database), or by deconstructing the chemical structure and calculating its formation energy using the component contribution method. Thereafter, the species involved in a reaction are combined (with consideration for stoichiometry) and the thermodynamic feasibility of the pathway is estimated by taking the sum of the reaction Gibbs free energy of each participating reaction. The node takes as input pathways in SBML format and returns annotated pathways (with thermodynamics information for each reaction, see Methods section for further details) also in SBML format.

**FBA**^i^ (Flux Balance Analysis) is used to calculate target production fluxes of the designed pathways. To perform FBA on a heterologous pathway, this tool first merges a heterologous pathway with a user-specified GEM model. This enables FBA to consider whole-cell conditions for the theoretical production of the user’s target molecule. The tool uses the CobraPy package to perform FBA^44^. The following native CobraPy methods are supported including FBA and parsimonious FBA (pFBA). The tool also contains an in-house developed method to consider the potential burden that the production of a target molecule may have on the cell and the impact of the target itself. We name the method “fraction of reaction”, and include the following steps. First the FBA for the biomass reaction is optimized and its value is saved as its optimum (note that by default the tool first optimizes to the biomass reaction, but the user may specify any reaction he so wishes). Then the upper and lower flux bounds of the biomass reaction is set to the same value, as a fraction of the optimum (default is 75%), and forces that flux to go through the biomass reaction regardless of the other set objective. Then the target reaction is optimized and the result of that flux is then reported. The node takes as input pathways like those produced by RP2Path and a strain model both in SBML format and returns annotated pathways (with calculated fluxes, see Methods section) in SBML format.

**Rank Pathways**^j^ ranks a given a set of heterologous pathways to reveal what are the most likely pathways to produce the target molecule in an organism of choice. It uses four different criteria: target product flux calculated by FBA, thermodynamic feasibility, length of the pathway, and reaction score based on enzyme availability calculated by RetroPath2.0. The weights are optimized by computing a global score for all pathways, ranking the collection, and optimizing weights such that the closest predicted pathway to any literature pathway for the same target is found on the top of the ranked list (for more information refer to section 2.3 and the Methods section). The node takes as input annotated pathways in SBML format and returns a ranked list of pathways also in SBML format.

**Selenzyme**^k,17^ is an open-source tool that performs enzyme sequence selection from a reaction query. The tool can be queried using a reaction template such as the reaction rules in RetroRules. This feature makes this tool especially useful in combination with RetroPath2.0. Selenzyme performs a reaction similarity search in the reference reaction database Metanetx^40^ and outputs the sequences annotated for the closest reactions. The tool provides several scores that can be combined in order to define an overall score. Scores are given for reaction similarity, conservation based on a multiple sequence alignment of the result, phylogenetic distance between source organism and host, and additional scores calculated from sequence properties. The Selenzyme node takes as input pathways in SBML format and returns annotated pathways (with UniProt ID for each reaction, see Method section) also in SBML format. A wrapper providing docker encapsulation for the Galaxy workflow is available^l^.

**SBML to SBOL converter**^m^ provides the mapping from the theoretical space to the practical space. The node takes a pathway model (encoded in SBML) as input, and returns a collection of placeholders for the subsequent design of the synthetic DNA that is required to encode the enzymes defined in the pathway model (encoded in SBOL). The converter first parses the SBML model, and extracts a user-specified number of homologous enzymes for each metabolic reaction. Synthetic gene design templates, in the form of SBOL *ComponentDefinitions*, are generated for each enzyme, each consisting of an (enzyme) coding region (specified by a Uniprot sequence identifier), 5’ and 3’ flanking regions for downstream assembly, and - optionally - ribosome binding sites of user-specified translation initiation rates, allowing for the control of translational regulation. The SBOL document contains no sequence data, but acts as a template to be passed onto the next node, PartsGenie.

**PartsGenie**^n^ is an established web application for the design of reusable synthetic DNA parts^45^. It supports the integrated design and optimisation of ribosome binding sites, coding sequences and other features, providing a multi-objective optimisation algorithm that simultaneously optimises translation initiation rate and codon usage along with elimination of repeating nucleotides and unwanted restriction sites. Furthermore, PartsGenie also implements guidelines from DNA manufacturers to optimise sequences for *synthesisability*, including the reduction of both local and global GC content. The PartsGenie node provides a wrapper for this functionality, taking in the “template” SBOL document from the preceding SBML to SBOL converter step as input, and using this a set of instructions for PartsGenie. The PartsGenie node then designs and optimises synthetic DNA sequences for each gene in the template, and updates the SBOL document with these novel sequences.

**OptDoE**^o^ combines selected genetic parts and enzyme variants for the desired. This node, based on the optimal design of experiments OptBioDes library^25^, accepts as input the pathways in SBML format annotated with the enzyme variants and the collection of genetic parts consisting of plasmid copy numbers of the vector backbone, resistance cassette, promoters, and terminator in SBOL format and registered in the SynBioHub repository. The *D*-optimal experimental design algorithm is based on a logistic regression analysis with an assumed linear model for the response evaluated based on its *D*-efficiency, which compares the design with an orthogonal design.

**DNA weaver**^p^ devises cloning strategies using either Golden Gate Assembly or Gibson Assembly to obtain plasmids for each combination of genetic parts selected by the OptDoE node. As both assembly methods have practical limitations, the algorithm first considers Golden Gate assembly using the type-2S enzymes BsmBI, BsaI, or BbsI (in this order) and defaults to Gibson Assembly, although this order of preference can be changed by the user. the resulting assembly strategies produce “scarless” plasmids whose sequence is the direct concatenation of the sequences of the plasmid’s parts. The node output is a spreadsheet featuring a list of all the primers required to extend the standard genetic parts with sequence homologies necessary for the assembly, and a list of all PCRs and fragment assembly operations required to obtain the desired plasmids. The assembly strategy is optimized to maximize primer reuse between constructs, and optimize assembly homologies, via the DNA Weaver framework^46^.

**LCR Genie**^q,47^ is a web-based tool for supporting the design of bridging oligos, which are required for annealing together individual synthetic DNA parts (designed by PartsGenie) into multi-gene plasmid assemblies, designed by OptDoE. The LCR Genie node provides a wrapper for this functionality, taking in an SBOL document containing numerous combinatorial plasmid assemblies, and designing bridging oligos necessary for assembly via the ligase cycling reaction method. The LCR Genie node performs analogous functionality to the DNA weaver node (supporting multi-part assembly but by a different experimental method) and as such, its output format matches that of DNA weaver.

**Pathway Visualizer**^r^ provides users an interactive web interface for exploring predicted pathways and their associated annotations. The tool is based on HTML and JavaScript only, which draws it as a “dependency-free” tool easy to set up locally for the user. Possible user interactions are pathway highlighting, cofactor handling, and the viewing of information at the levels of pathways, reactions, and involved compounds. The node takes as input pathways in SBML format.

The Galaxy-SynBioCAD portal does not currently support the visualization of SBOL files such as those produced by PartsGenie and OptDoE, however, these files can be downloaded and can easily be visualized using online tools such as Visbol^s^.

The Galaxy-SynBioCAD portal also supports other nodes not listed above that perform simple operations like uploading a file, extracting taxonomy ID, or native metabolites from a GEM SBML file. All these nodes are fully described in the Supplementary Information and the Node Documentation file found on the portal.

### Building workflows with nodes

As described in the above section, the SynBioCAD-Galaxy portal contains a collection of tools that have simple standardized input and outputs. These “nodes” are well documented and intended to perform a single well-defined task. To create more complex tasks, these tools may be chained together. We present three exemplar workflows.

#### Retrosynthesis and Pathway Enumeration

The workflow (Figure 1) generates theoretical possible pathways for the production of a target molecule in an organism of choice. Three key steps are performed in this workflow. First, using the RetroPath2.0 node, it generates feasible metabolic routes between a collection of chemical species contained within a GEM SBML file of the selected organism, a target molecule that the user wishes to produce, and reactions rules extracted from RetroRules. That metabolic network is then deconstructed into individual pathways using the RP2paths node. Lastly, those individual metabolic pathways are converted to SBML files using the *Pathways to SBML* and *Complete Reactions* nodes. The former generates SBML files describing the individual heterologous pathways while the later adds the appropriate cofactors and removes duplicate pathways.

**Figure 1.**
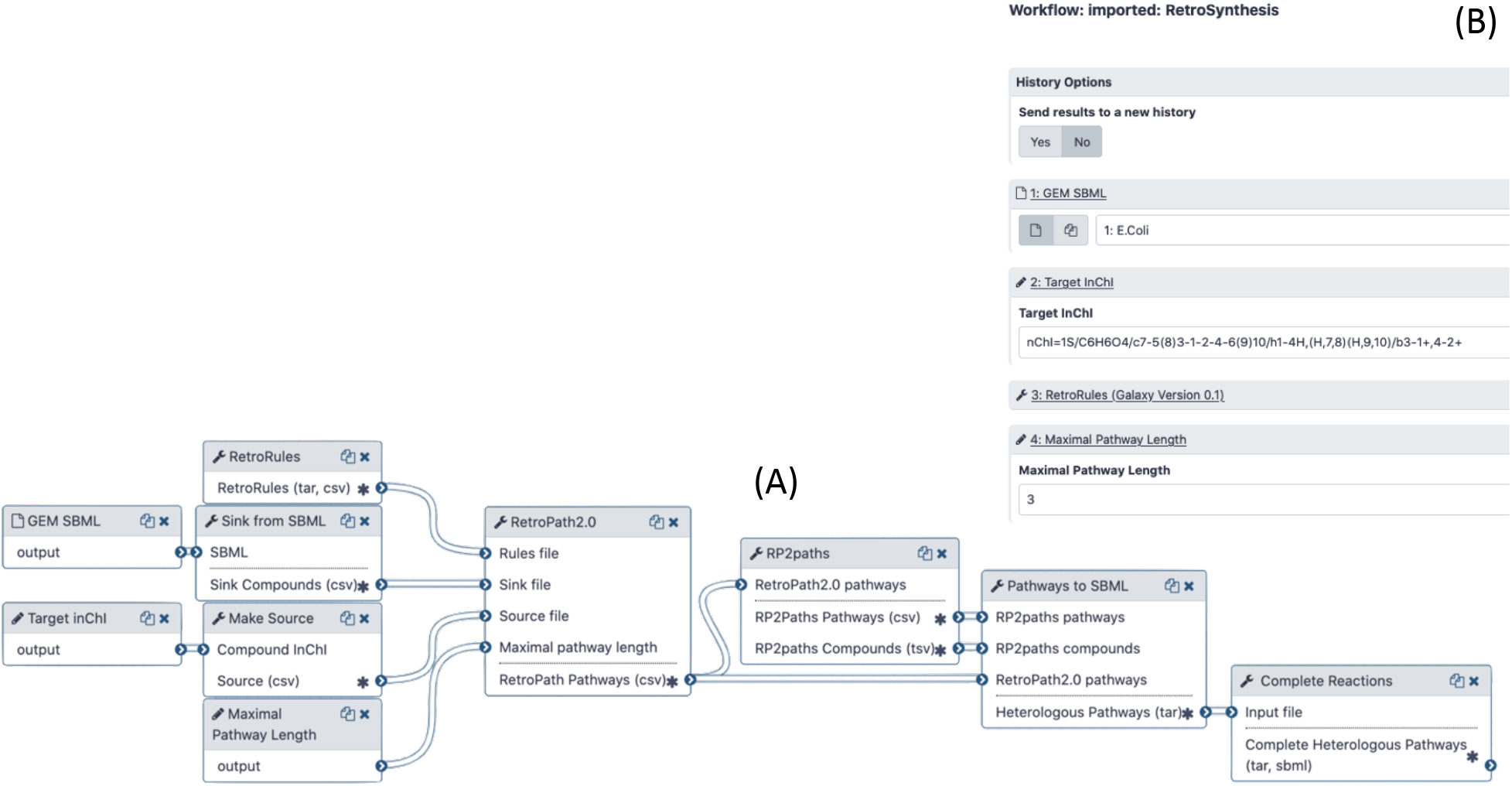
The RetroSynthesis workflow as seen in the Galaxy portal. (A) The workflow in workflow editor. (B) The workflow menu upon executing it. There, the user must specify the GEM SBML model of the host organism, the InChI structure of the target molecule, and the maximal pathway length. The workflow generates a collection of heterologous pathways for the production of the target into distinct SBML files.

#### Pathway analysis and ranking

Given a set of pathways generated by RetroPath2.0, this workflow informs the user as to the theoretically best performing ones based on the four criteria mentioned on the previous section (node Rank Pathway: target product flux calculated by FBA, thermodynamic feasibility, length of the pathway, and reaction score based on enzyme availability). In the previous workflow, molecules contained within a full SBML model are used to compute heterologous pathways. As a result, the calculated heterologous pathways can easily be merged into the full organism model, enabling whole-cell context to calculate the potential flux of a given target. Under such simulation conditions, the analysis that returns a low flux may be caused by the starting compound itself not having a high flux, or the cofactors required having a low flux, while the pathways with high flux would be caused by both the starting compound and the cofactors being in abundance. In either case, bottlenecks that limit the flux of the pathway may be identified and pathways that do not theoretically generate high yields can be filtered out. Furthermore, the production of heterologous molecules in an organism often causes a burden on the growth of the cell. To emulate such a condition, we use here the method named “*fraction of reaction*” and described in the previous section for the FBA node. The method forces a fraction of its maximal flux through the biomass reaction while optimizing for the target molecule. The reaction score that probes enzyme availability for the chemical transformation is also taken into consideration, where high values favour less promiscuous reaction rules and express better confidence. Finally, the length of the pathway is taken into consideration as well, here shorter pathways are favored over longer pathways. The results of the pathway analysis are combined using a weighted mean, and a single global score is computed. The results may be graphically inspected by the user using a Galaxy embedded visualizer that displays the heterologous metabolic routes for the production of a target molecule in an organism of choice, where complete descriptions of the chemical species, reaction and pathways are displayed (Figure 2).

**Figure 2.**
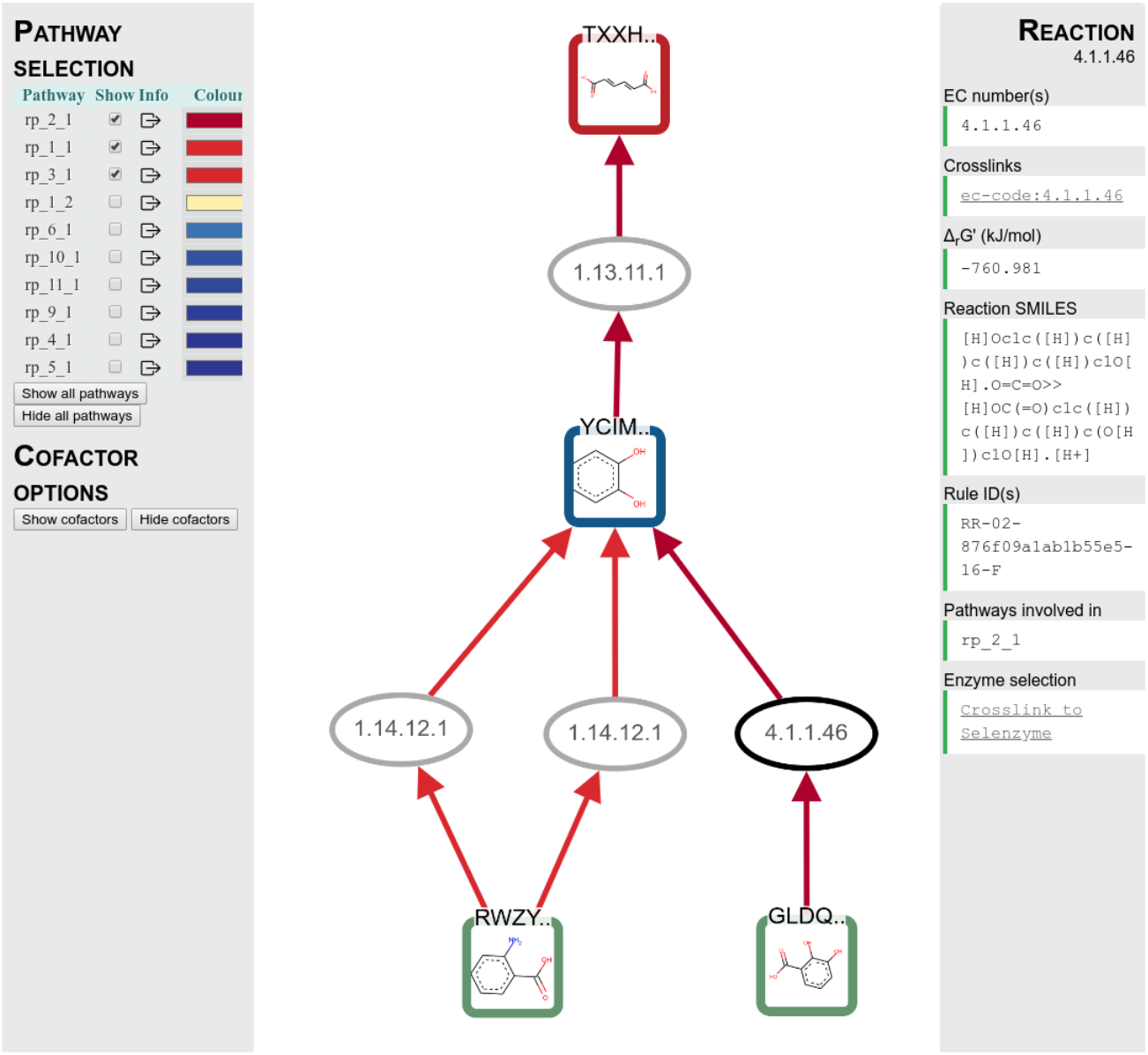
Top scored enumerated pathways. The example plots the top 3 ranking pathways for the production of muconic acid in E. coli after running the “Pathway Ranker” workflow as seen on the portal Visualiser. The squares depict the molecules and the ovals the reactions. The green squares are the compounds that exist in the GEM model, the blue squares are intermediate compounds, and the red square is the target. Each compound and reaction can be selected and the right hand side displays details of the selection (here for the reaction 4.1.1.46). The panel offers also a link to look up the reaction on the Selenzyme web service to manually search enzymes that may perform such chemical transformations. On the left hand side is the ranked list of pathways predicted, color coded so that the best theoretical performing ones have warmer colors. The user may inspect the pathway as a whole by selecting the boxed arrow. This action displays on the right hand panel information on the pathway including the number of steps, its thermodynamic feasibility, its flux and its global score. The user can also display the cofactors for all the reactions by selecting the “Show cofactors” button on the left side panel.

#### Genetic Design

This workflow encodes the top-ranking predicted pathways from the previous workflow into plasmids intended to be expressed in the specified organism. First, the Selenzyme node is executed to return a user defined number of UniProt ID’s associated with each reaction. Then a maximum number of pathways, as defined by the user, are converted to SBOL. The next tool, PartsGenie, then retrieves the DNA sequences of the predicted enzymes based on their Uniprot ID, performs a codon optimization and creates a first level of library based on those, adding before the CDS some specific strength calculated RBS. These constructions are then used by OptDoE to generate a defined size library of plasmids, expressing at various levels the genes coding for the multiple enzymes present in the predicted pathways. The other genetic parts required by this software (origin of replications, promoters, terminators and markers) are either provided by a default list or a specific list of parts provided by the user which needs to refer to parts stored in SynBioHub. The Galaxy tool “OptDoE Parts Reference Generator” has been written for that purpose. This final library is generated in a SBOL format and can then be used as an input to other softwares or visualised using tools implementing the SBOL visual standard. The Genetic Design workflow ends with two different tools tackling the library construction problematic: LCR Genie that propose an assembly strategy using the Ligase Chain Reaction method and DNA weaver that calculate the optimal synthesis plan and the assembly protocol following either a Golden Gate or a Gibson Assembly method. The output of LCR Genie or DNA weaver are excel files containing the full sequence of the plasmid library and of the intermediate parts required to construct them.

### Benchmarking with literature data

Although criteria like target product flux, thermodynamic feasibility, pathway length, and reaction score based on enzyme availability inform the user as to the best potential candidate pathway to produce a compound of interest, we are interested in ranking pathways combining these criteria in such a way that a global score value may be used to determine what are the best candidates.

To achieve this, a list of experimentally expressed compounds in engineered organisms (*E. coli, S. cerevisiae, B. subtilis and Y. lipolytica*) reported in the literature was collected. For each pathway and each heterologous pathway reaction we compiled the EC number of the reaction along with the substrates and products of the reaction. This list may be found in the Supplementary Information. Each target compound within that list was used to run the “*Pathways Analysis and Ranking*” workflow described in the previous section to generate a collection of predicted pathways that produce the same target molecule in the same host organism than those reported in the literature (Figure 4.A). Following that, the predicted collection of pathways were compared with their corresponding literature pathways using a matching algorithm described in the Methods section and illustrated in Figure 4.B. The predicted pathway with the highest similarity (and above a given similarity threshold, see Supplementary Information) was flagged as the best performing pathway among the collection.

**Figure 3.**
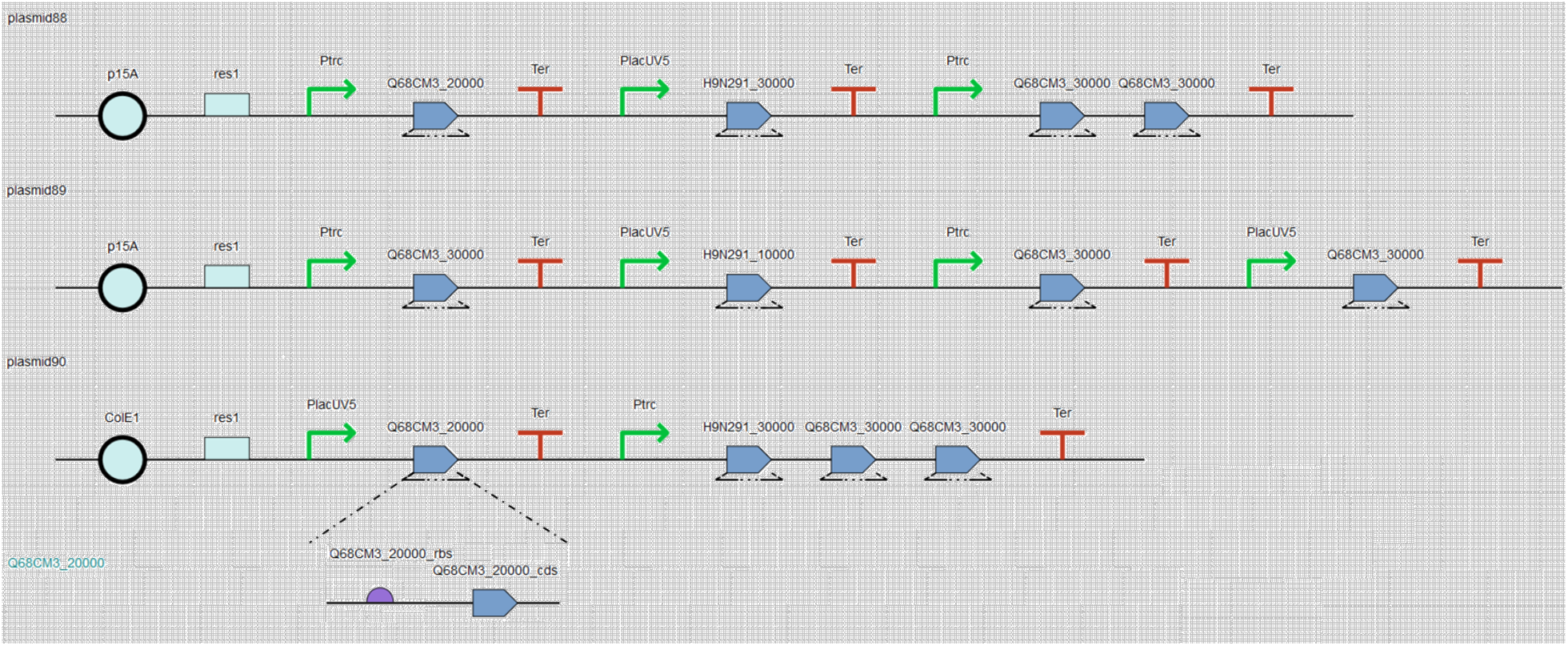
Architecture of the constructed library of plasmids in SBOL format. The figure illustrates three layouts in SBOL format representing each one of the plasmids implementing the heterologous pathways producing muconic acid in E. coli. The SBOL pathway layouts are visualized using the web service visBOL.

**Figure 4.**
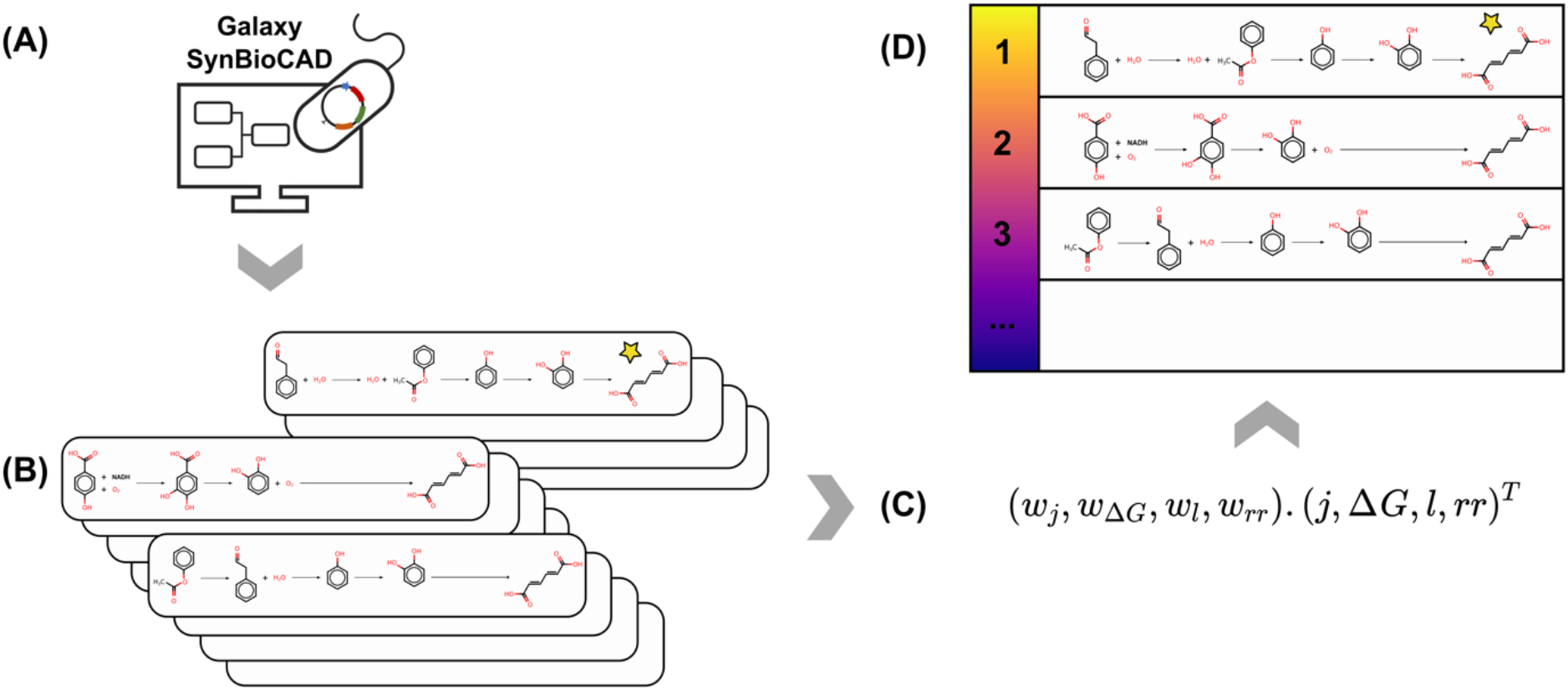
Scoring workflow predicted pathways with literature pathways. (A) The RetroSynthesis and Pathway Analysis workflows are run on a given literature target molecule (here muconic acid) to generate a collection of predicted pathways. (B) Comparison is performed between all the predicted results to the literature pathway (marked with a star). (C) The weights associated with the different criteria of the predicted pathways are combined using a weighted mean to calculate the global scores of the pathways, where: j is the target product FBA flux results, ΔG^m^ is the thermodynamics result, l is the length of the pathway, rr is the enzyme availability reaction score, and ware the weight parameters. (D) The global scores are then used to rank the list. This process is repeated for every literature pathway and the weights are optimised such that a maximum of literature pathways are found on the top of the ranked list.

In the current study, we assumed that the literature pathway is among the best performing predicted pathways, but we also hypothesize that other predicted pathways may be valid as well. The similarity with literature pathway is thus used to determine the importance (weights) of the criteria contributing to the scoring of the pathways and generate a global score to rank the predicted pathways from best to worst (Figure 4.C).

In order to find the literature pathway in the top scored predicted pathways we used the ranked-biased overlap algorithm^48^ to score the performance of the weights (see Method section). The algorithm produced the following list of optimized weights: pathway length weight: 0.73%, reaction score weight: 12.46%, FBA target product flux weight: 32.4%, and thermodynamics weight: 54.4%.

Using the above weights, Figure 5 shows the results of the ranked-biased overlap optimization schema. Each row is a ranked list of collections of predicted pathways for a given target molecule, where on the left-hand side are the best ranking pathways. The color code shows the global score that was used to rank the pathways. The black boxes correspond to the literature pathways that are the most closely similar to the literature pathway (see Supplementary Table SX for score values of literature pathways). Overall, we find that our “*Pathways Analysis and Ranking*” workflow after adjusting the weights using ranked-biased overlap has a 65.4% success rate (53 out of a total of 81) in retrieving the literature pathway among the top 10 predicted pathways.

**Figure 5.**
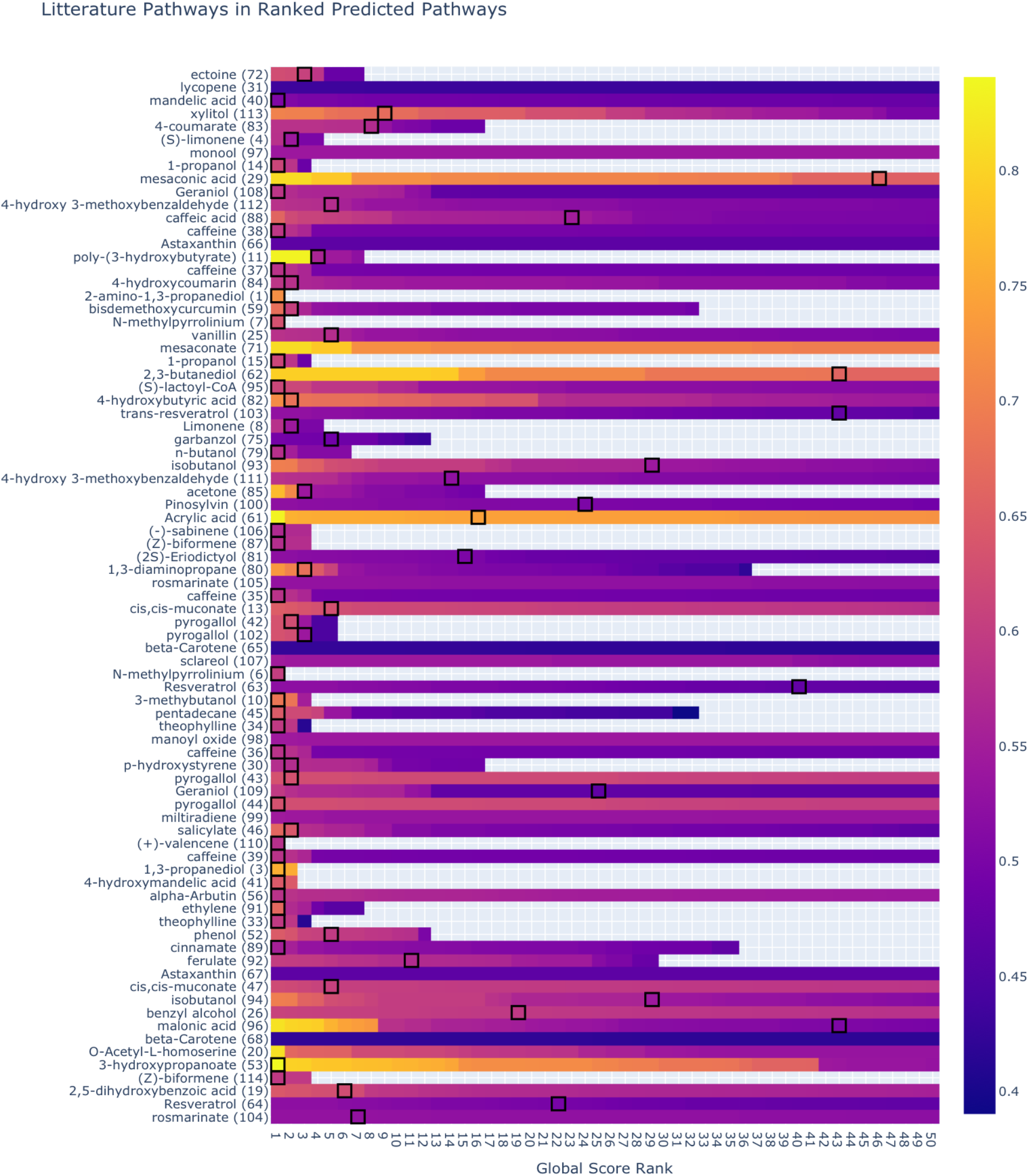
Results of the optimization showing the first 50 ranked predicted pathways for the predicted pathways. The black boxes show the location of the closest predicted pathway to the literature pathway. The optimization algorithm balances the weights of the criteria of pathways such that the literature pathways are on the top rank of the predicted pathways. If a row does not contain a black box then the identified literature pathway is not found within the first 50 predicted pathways.

## Discussion

We have presented in this paper several Galaxy workflows to design pathways in host organisms. These workflows have been built using 20 different computational tools (named nodes) currently present on the platform (*cf.* section 2.1 and Supplementary Information for a complete list of nodes). Chaining the nodes together to form workflows was made possible only because the input and output of each node were standardized. As far as standardization is concerned, we chose community adopted standards like InChI and SMARTS for compounds and reactions, SBML for pathways and strains, and SBOL for genetic constructs. Considering all the workflows that could potentially be created on Galaxy SynBioCAD with the current nodes, the end-to-end process offered in the portal starts from the specifications of the targeted compounds and the selected hosts, to the DNA parts to be synthetized depending on the assembly protocol (LCR, Golden Gate, Gibson are currently offered). Combinatorial layouts of the pathways can also be generated via the OptDoE node.

The pathways generated by the workflows have been compared with literature pathways, and in order to maximize the number of times the literature pathways were found in the top pathway list returned by the workflows, a ranking function was developed (see section 2.3). That function is a weighted sum of four criteria: target product flux, reaction thermodynamic feasibility, reaction score based on enzyme availability, and pathway length. Interestingly, the pathway length weight alone seems to be a bad predictor of the quality of the pathways. We suspect that the reason this criterion has such a meager influence on the global score stems from the fact that, for a given target molecule, we have most often only identified a single pathway that describes its production in the literature. Therefore, while our workflow returns a plethora of heterologous pathway solutions to produce a given target (some of which are shorter than the literature reported) the scoring method penalizes the shorter metabolic pathways that might otherwise be considered to be better solutions. For a better optimization solution regarding that parameter, we would need to compare multiple pathways that produce the same target with different lengths and favor shorter length pathways. Experimental validation would be needed to confirm that shorter pathways are better predictors, which is out of the scope in the current study. For the time being, lack of such data in the literature leads the length to have a small influence on the global score.

While searching literature pathways in the set of pathways produced by our workflows is appropriate, this does not mean other pathways generated by the workflows are not valid and cannot be engineered. In order to assess the validity of all predicted pathways, one strategy that has been used for synthesis planning in chemistry is the double-blind testing strategy performed by a pool of participants^49^. In that strategy neither the participants nor the conductors are aware of the origin of the pathways, and the participants are asked to flag pathways they deemed valid without having explicit information on pathways found in the literature. Such a method could be applied here to further refine the ranking function.

To summarize, the Galaxy-SynBioCAD portal proposes the first set of synthetic biology computational tools in a Galaxy framework^35^. We chose Galaxy as our workflow system because the tools found in the ToolShed^36^, have reached way beyond genome analysis for which Galaxy was originally developed. Just by focusing on tool categories found relevant to the present manuscript, one can cite proteomics, transcriptomics, metabolomics, flow cytometry analysis, and computational chemistry. Several communities are using Galaxy and many papers can be found online^50^ for microbiome (267 items are found as of 07/04/2020), plants (258 items), diseases like cancer (312 items), and drug design and discovery (75 items). However, the library hardly contains references related to biotechnology (4 items) and even fewer to synthetic biology and metabolic engineering.

The current offering in Galaxy-SynBioCAD focuses on providing tools for pathway design. However, as Galaxy-SynBioCAD is a community effort, we anticipate our tool set will grow. Regarding pathway design tools, many of the software products listed in the introduction could be considered to be added to the portal. In particular, strain design including knockout genes to maximize targeted product fluxes, could easily be implemented via the FBA tools making use of Cobrapy (see section 2.1). Additionally, there are already Galaxy workflows to take up and analyze metabolomics flow cytometry data in ToolShed^36^, and these workflows could directly be incorporated into the portal to deal with data generated in the ‘Test’ step of the DBTL cycle. As mentioned in the introduction several open source software products deposited in GitHub^26–29^ could cover the ‘Build’ step and eventually provide drivers to automated constructions. Regarding the ‘Learn’ step in DBTL, the OptDoE tool (cf. section 2.1) could easily be adapted to propose new designs as it was done in Carbonell *et al*., more complex approaches to be considered are methods that make use of machine learning as in Borkowski *et al*.^51^. While all design examples provided in the current paper are for engineering pathways in host organisms, because of the recent development of models (similar to GEM models) for cell-free systems^52^, one can also consider adapting the portal for design and engineering in cell free.

All of the above suggested additions could be implemented in our portal with relatively small efforts. There are other applications that could be envisioned beyond pathway design and engineering. For instance, as shown in Delepine *et al*.^10^ retrosynthesis software can easily be adapted to design biosensors, and tools used in Cello^3^ or Pandi *et al*.^53^ that respectively propose designs for genetic logic circuits and metabolic neural network biocomputation could also be considered.

## Methods

### Pathway annotation

Some results generated by the workflow nodes produced in this study cannot be readily stored in the SBML files natively (example: reaction rule, thermodynamics, etc…). As such, we elected to enrich the SBML format in such a way that our information can be stored directly within the SBML file without breaking any standard of the original file. Because SBML files are based on XML, new XML annotations are created that are outside the standard scope of a SBML file and thus are ignored by any standard SBML readers^5^. We denote that enriched file format rpSBML and is compatible with any other SBML readers (additional details can be found in Supplementary Information).

Standard SBML extensions are also used in this project. The “groups” package is used to link the heterologous reactions and chemical species to identify them easily, as well as classifying the chemical species that are main actors in a heterologous pathway^54^. While the FBC package is used to define the FBA simulation conditions^55^.

### Literature Pathways matching algorithm

An important requirement in this project is the need to compare two different metabolic pathways, and quantify the degree of similarity between the two. This is used when searching if a literature pathway can be found in a list of pathways produced by our workflows. To this end we wrote a matching algorithm that compares SBML files and calculates a similarity value using the following criteria:

- Chemical species

○ Chemical structure (InChiKey)
○ Public database cross-references (MIRIAM)
- Reactions

○ Substrates/Products similarity
○ EC number
- Pathway

○ Length

A more complete description of the algorithm may be found in Supplementary Information. In short, all the chemical species between two SBML pathways are first compared and the best matching ones are coupled ensuring 1:1 matches. The species match is then used, when comparing two reactions, to match all the substrates and products between two reactions. The EC number is also used as a criterion to determine the similarity of two reactions. If two reaction EC numbers have the same main class, subclass and sub-subclass then these reactions are considered to be the same. The full EC number is however not ignored, as two reactions with exactly the same EC number would score higher than two that have up to the third similar EC numbers. Finally, a penalty score is applied to the similarity value if the length of the pathways differ. The output is a single similarity value that can be used to determine what is the closest predicted pathway to the literature reported pathway.

### Ranked-Biased overlap method

To validate predicted pathways, we use literature reports of engineered pathways and compare the results corresponding to the same target compounds. The underlying assumption we are making is that the literature pathway must be among the best performing pathways, but must not necessarily be the best performing one.

Our workflows generate a collection of heterologous pathways for the production of a given compound. The first step involves computing a similarity value for every predicted pathway with a given literature pathway and selecting the top ranking one as the member that best matches the litterature pathway. Thereafter, a global score is computed (for a given set of weights associated with the criteria of the pathway, see section 2.3.) that also returns a ranked list with the better ranking pathways on the top of the list. For this ranked list, we must determine if the corresponding litterature pathway is on the top or not. To this end, we use the Rank-Biased overlap algorithm (RBO)^48^.

This algorithm offers a few advantages that particularly suit our needs. First, it can compare lists of disparate sizes. Indeed, during our optimisation, we select only the best or the closely matching similar predicted pathways to the literature pathway as the best performing pathways. Secondly, RBO provides a parameter to control the degree of importance of matches on the top of the list (also called top-weight). In other words, it controls the sharpness of increase in the score as the literature pathway finds itself on the top of the ranked list of pathways. We use this parameter to loosen the requirement of the literature pathway to be necessarily on the absolute top of the ranked list, while still giving a scoring advantage that the pathway finds itself in a better position (see Supplementary Information).

### IT Architecture

The Tools in Galaxy-SynBioCAD can be used in a stand-alone way or chained into workflows. The source code of each tool is open-source and available in GitHub repositories^t^. The IT architecture of SynBioCAD is based on the Docker^u^ framework. Each component of this architecture is running with Docker, from web interface to network management.

Each container is confined into its own network so that it is not reachable by any other component. To make communications possible between two containers, one of them is put into the other’s network by choosing the most secure option. As an example, in order to enable communication between the Galaxy container and a RESTful tool, we have two options: (1) put the tool into the Galaxy container’s network or (2) puts the Galaxy container into the tool’s container. Let’s say we have several RESTful tools, the first option stores all tools into the Galaxy container’s network that enables the communication between tools themselves; which also breaks the network confinement and decreases the global system’s security level. The second option preserves the network confinement between tools while making possible communications between each tool and the Galaxy container. In the SynBioCAD framework, we choose the option that optimizes the security for all containers that need to communicate (reverse-proxy/Galaxy, Galaxy/database…).

The overall architecture is illustrated in Figure 6 details regarding Galaxy and tool services are detailed next and an example is provided for RetroPath2.0.

**Figure 6.**
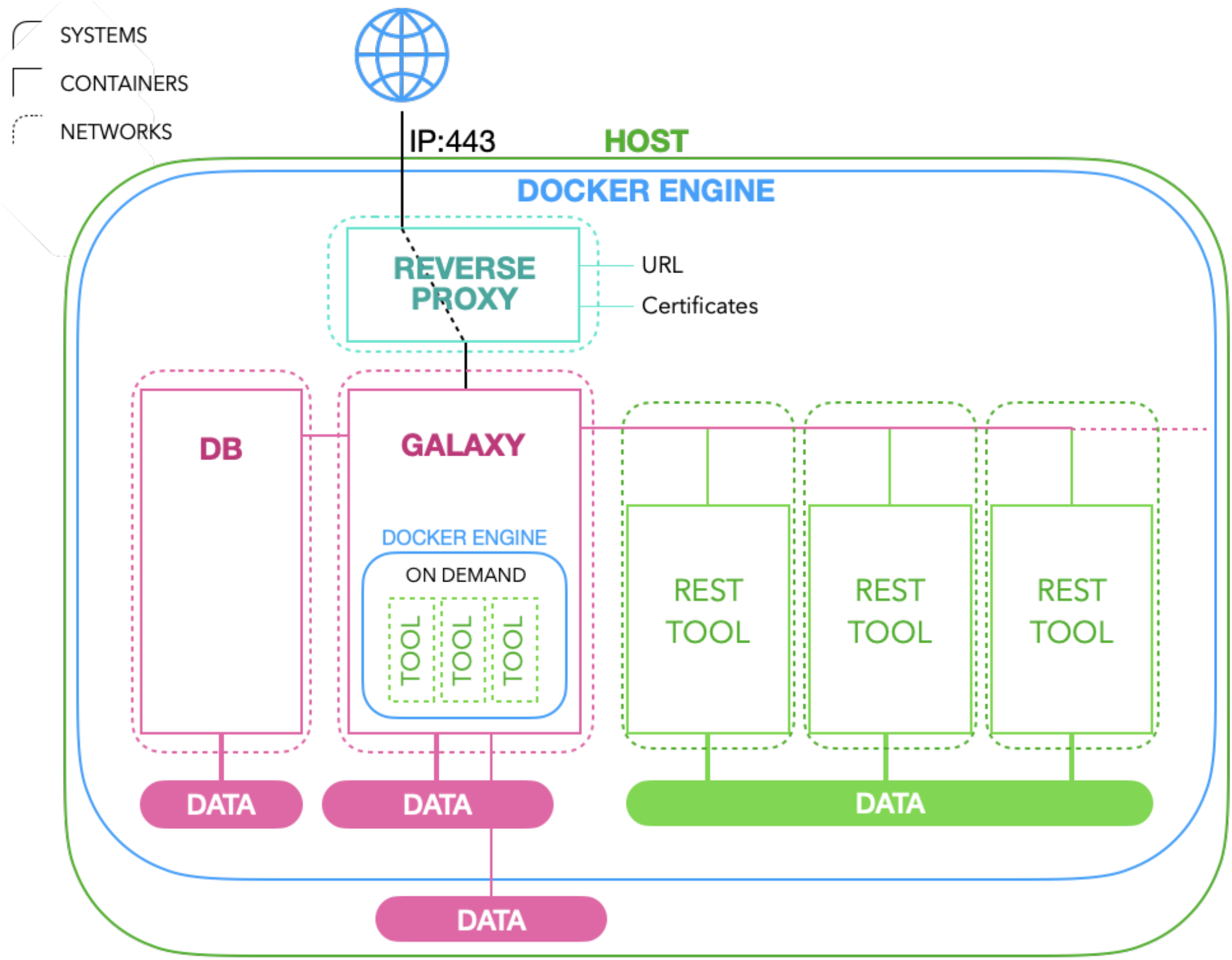
SynBioCAD IT architecture overview. Each component is embedded in the Docker environment installed on the (virtual or physical) host. In addition, each container is confined into its own network and therefore is unable to be reached by any other components. To make communication possible between two containers, we put them into the same network by preserving networking confinement.

#### Galaxy Service

The main brick is the Galaxy service which is the web interface for end-users and orchestrates tool executions. The Docker Galaxy system is based on the following bricks: (1) Install container runs once and downloads galaxy project sources from the web^v^. (2) Galaxy container downloads from the web the Galaxy project, and runs it within the galaxy image^w^. This step is time-consuming for the first time due to Galaxy’s initialization. (3) Database containers are used by Galaxy for users accounts and dynamic web pages. This service is dedicated to Galaxy service. About data storage, each container described above relies on Docker data volumes for storing persistent data. Concerning networking level, the database container is confined into its own network so that, by default, it is not reachable by any other container. The Galaxy container is confined into its own network so that, by default, it is not reachable by any other container. However, this service is also part of the database service network in order to communicate with it. All these bricks are orchestrated by the Docker Compose tool and embedded in a docker-compose file.

#### Tools Services

All tools available in the Galaxy-SynBioCAD portal are dockerized and run in two different modes: (1) on-demand where tool images are instantiated each time the tool is requested. These tools run within the Galaxy container, which embeds a Docker engine. (2) RESTful, where a REST service is always up and embeds the tool. These services run next to the Galaxy container and communicate with the Galaxy container through Docker networking. Tools available in Galaxy have to be deployed within the Galaxy container.

###### RetroPath2.0 Service Example

RetroPath2.0 runs as a REST service and is deployed next to the Galaxy container. Its deployment is based on two containers.

Install container runs once and downloads data (e.g. RetroRules) into the data volume. REST container is a RESTful container that embeds the tool (through the tool image) and waits for requests.

Concerning data storage and networking level, RetroPath2.0 follows the policy described above. All sources can be found on GitHub and are splitted into two different repositories:

1. Docker image embeds all packages needed for running the tool. In addition, RetroRules data is downloaded within the image and the KNIME Analytics portal is installed.
2. Galaxy wrapper contains the necessary code for displaying the tool web page and to trigger the tool algorithm.

## Acknowledgements

MdL and JLF acknowledge funding provided by the infrastructure IBISBA (Horizon 2020 under grant agreement No 730976), TD and JLF funding provided by BioRoBoost (Horizon 2020 under Grant agreement No 820699) and PC, NS and JLF funding from the Biotechnology and Biological Sciences Research Council (BBSRC) and the Engineering and Physical Sciences Research Council (EPSRC) under grant ‘Centre for synthetic biology of fine and specialty chemicals (SYNBIOCHEM)’ (BB/M017702/1). NS acknowledges further funding from the BBSRC under grant ‘GeneORator: a novel and high-throughput method for the synthetic biology-based improvement of any enzyme’ (BB/S004955/1) and from the University of Liverpool. PC also acknowledges support from the Universitat Politècnica de València Talento Programme. VZ was supported for this work by The Edinburgh Genome Foundry funded by the BBSRC (BB/M025659/1, BB/M025640/1, and BB/M00029X/1 to Susan Rosser) and the BBSRC/MRC/EPSRC funded UK Centre for Mammalian Synthetic Biology (BB/M0101804/1 to Susan Rosser) as part of the RCUK’s Synthetic Biology for Growth programme.

## Authors information

JLF designed the study and wrote the main text of the paper. MdL dockerized and integrated all the Galaxy-SynBioCAD tools into Galaxy nodes, he also performed the literature benchmarking and wrote the corresponding section along with the supplementary information. NS, PC, TD, ZA integrated several tools in the portal with the help of MdL and wrote the corresponding description. JH supervised code development and dockerization for in-house tools. In addition, JH is in charge of deployment of all tools available in SynBioCAD web portal as well as the Galaxy platform itself (web, db, tools, networking). LF, FS, MM, PS tested and documented nodes, created workflows and wrote the corresponding description, and compiled literature pathways data.

## Footnotes

a galaxy-synbiocad.org

b github.com/Galaxy-SynBioCAD/RetroRules_image, github.com/Galaxy-SynBioCAD/RetroRules

c retrorules.org

d github.com/Galaxy-SynBioCAD/RetroPath2_image, github.com/Galaxy-SynBioCAD/RetroPath2

e github.com/Galaxy-SynBioCAD/rp2paths_image, github.com/Galaxy-SynBioCAD/rp2paths

f github.com/Galaxy-SynBioCAD/rpReader_image, github.com/Galaxy-SynBioCAD/rpReader

g github.com/Galaxy-SynBioCAD/rpCofactors_image, github.com/Galaxy-SynBioCAD/rpCofactors

h github.com/Galaxy-SynBioCAD/rpThermo_image, github.com/Galaxy-SynBioCAD/rpThermo

i github.com/Galaxy-SynBioCAD/rpFBA_image, github.com/Galaxy-SynBioCAD/rpFBA

j github.com/Galaxy-SynBioCAD/rpRanker_image, github.com/Galaxy-SynBioCAD/rpRanker

k github.com/synbiochem/selenzyme

l github.com/Galaxy-SynBioCAD/rpSelenzyme_image, github.com/Galaxy-SynBioCAD/rpSelenzyme

m github.com/Galaxy-SynBioCAD/rpSBMLtoSBOL_image, github.com/Galaxy-SynBioCAD/rpSBMLtoSBOL

n github.com/Galaxy-SynBioCAD/PartsGenie_image, github.com/Galaxy-SynBioCAD/PartsGenie

o github.com/pablocarb/doebase, github.com/Galaxy-SynBioCAD/rpOptBioDes_image, github.com/Galaxy-SynBioCAD/rpOptBioDes

p github.com/Galaxy-SynBioCAD/DNAWeaver_image, github.com/Galaxy-SynBioCAD/DNAWeaver_image,

q github.com/Galaxy-SynBioCAD/LCRGenie_image, github.com/Galaxy-SynBioCAD/LCRGenie_image, github.com/Galaxy-SynBioCAD/LCRGenie

r github.com/Galaxy-SynBioCAD/rpVisualiser_image, github.com/Galaxy-SynBioCAD/rpVisualiser

s visbol.org

t github.com/Galaxy-SynBioCAD, github.com/brsynth

u www.docker.com

v github.com/galaxyproject/galaxy

w github.com/brsynth/galaxy_image, github.com/brsynth/galaxy-dind_image

