## Supplementary text for "Galaxy-SynBioCAD: Synthetic Biology Design Automation tools in Galaxy workflows"

**2 Enriched Metabolic Model File: rpSBML** **7**

**3 FBA: Fraction of Reaction** **7**

**4 Pathway Matching Algorithm** **8**

**6 Optimisation** **15**

### 1 Workflow Nodes

Here we list the utilities tools that are not described in the main paper. These are tools that help the user run the main nodes of the Galaxy platform.

#### 1.1 Sink from SBML

This node takes for input an SBML file, parses all the species within a user specified compartment, and uses the MIRIAM annotations to identify the species to retrieve their InChI structure description. The results are written to a RetroPath2.0 friendly CSV file format. In the advanced options one can specify the compartment from which the tool will extract the species from. The default is MNXC3, the MetaNetX code for the cytoplasm. If the user wishes to upload an SBML file from another source, then this value must be changed. Furthermore, FVA may be performed to remove all the dead-end metabolites from the list.

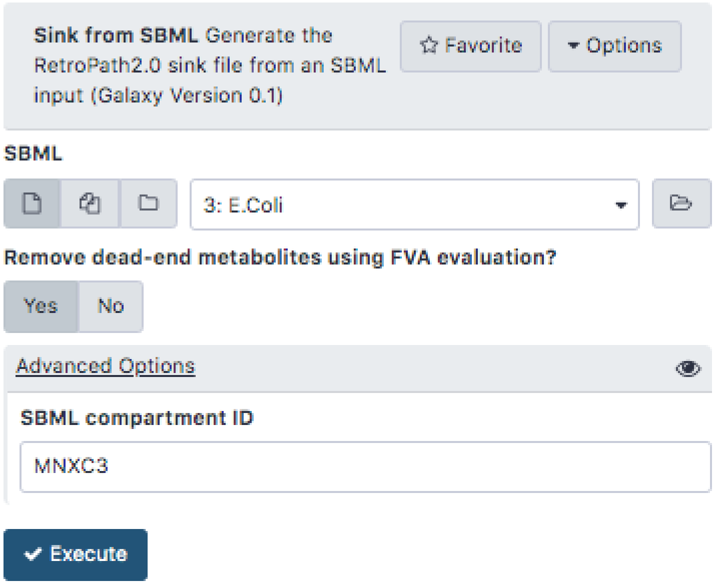

**Figure S1:** Sink from SBML node as shown in Galaxy. The user must specify a GEM SBML file and the compartment where the heterologous pathway will be expressed in and the tool returns all the chemical species contained within it. The user can also specify if the tool runs FVA to remove among the above list dead-end metabolites.

#### 1.2 Make Source

The node takes for input an InChI string and a compound name, and outputs a CSV file.

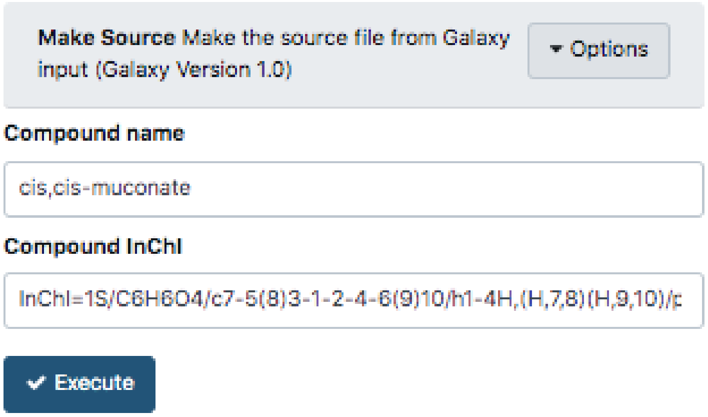

**Figure S2:** Make source Galaxy node. The tool takes a compound name and an InChI structure. The tool defaults, as shown here, are only for illustration purposes.

#### 1.3 Report

This tool takes as input an SBML or a collection in the form of a TAR and extracts the results of the analysis (FBA, thermodynamics, etc...), and outputs a CSV file that summarises the results.

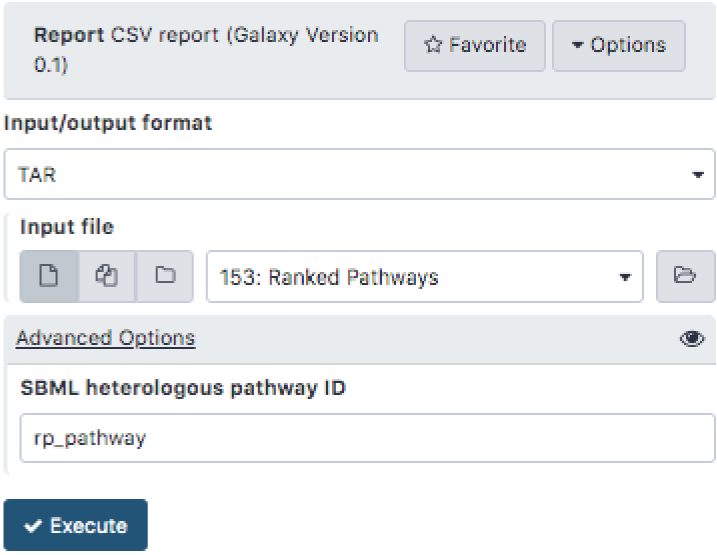

**Figure S3:** Report Galaxy node. The user specifies the collection of SBML files that are generated through the platform and the tool outputs a CSV that show the enriched information contained within the SBML file.

#### 1.4 Merge SBML

This node takes two different SBML files (or a collection in the form of a TAR) and merges one into another. Common usage of this tool includes merging a predicted heterologous pathway into a GEM model for further analysis (see the FBA tool).

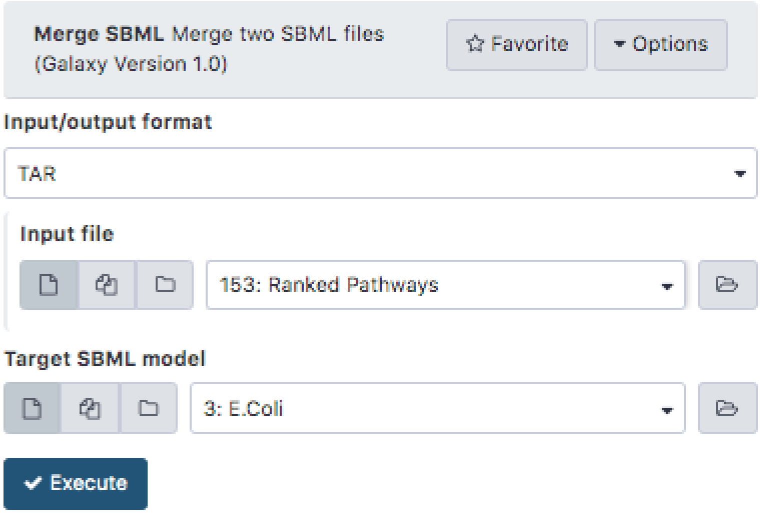

**Figure S4:** Merge SBML tool as seen on the Galaxy platform as a user is executing it. The user must specify the source model (either a single or a collection in the form of a SBML) and a target model in the source model will be added to.

#### 1.5 Extract Taxonomy

This tool extracts the NCBI taxonomy ID from a GEM SBML model. This tool is to be used with OptDoE or Selenzyme, where both need that information.

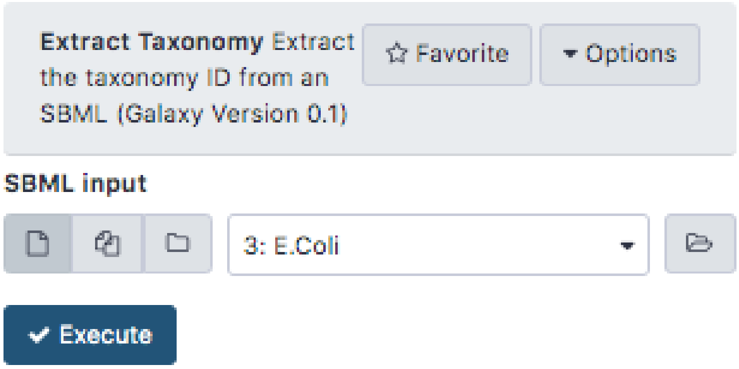

**Figure S5:** Extract Taxonomy tool that attempts to find the appropriate taxonomy ID from a GEM SBML file.

#### 1.6 OptDoE Parts Reference Generator

To run OptDoE, a list of genetic parts is required to calculate the possible combinatorial genetic designs for a given heterologous pathway. If none are provided, then the OptDoE tool provides a default list of parts, as shown in figure SX. However, because it is of particular interest to have the user provide his own genetic parts, this tool aids a user at creating a valid CSV with user defined genetic parts. For each part, the user must provide a name, the type of part that may include: Promoter, Origin, Resistance and Terminator and lastly the SynBioHub IP address.

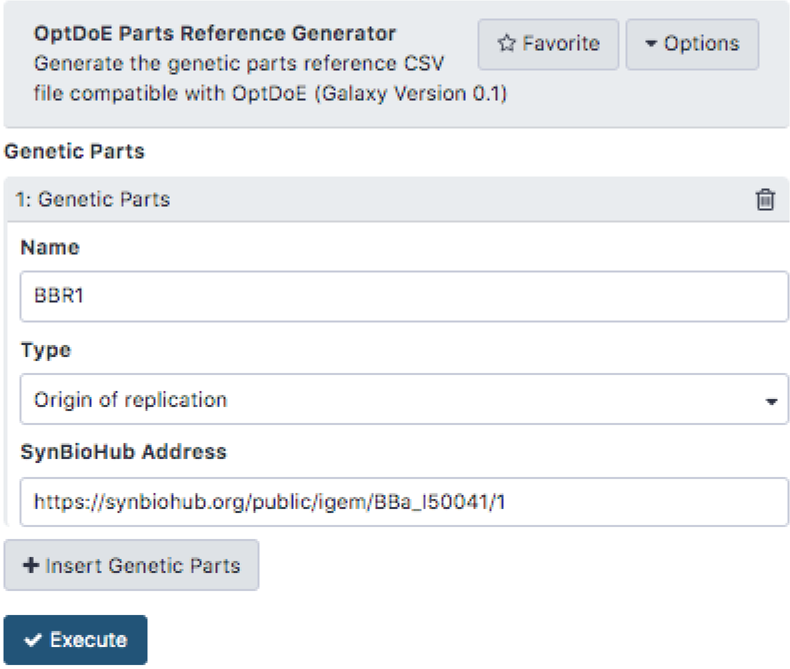

**Figure S6:** OptDoE Parts Reference Generator as seen on the Galaxy platform. The user can add as many genetic parts as he wishes by selecting the + Insert Genetic Part. For each, the user must specify the name, select the type of the part from a list that include: promoter, origin of replication, resistance and terminator. Last, the user must specify the SynBioHub address of the genetic part.

| Name | Type | SynBioHub |
|---|---|---|
| PlacUV5 | Promoter | https://synbiohub.org/public/igem/BBa_K1847014/1 |
| Ptrc | Promoter | https://synbiohub.org/public/igem/BBa_J56012/1 |
| BBR1 | Origin | https://synbiohub.org/public/igem/BBa_I50041/1 |
| p15A | Origin | https://synbiohub.org/public/igem/BBa_I50032/1 |
| ColE1 | Origin | https://synbiohub.org/public/igem/BBa_J64101/1 |
| res1 | Resistance | https://synbiohub.org/public/igem/BBa_I13800/1 |
| Ter | Terminator | https://synbiohub.org/public/igem/BBa_B1006/1 |

**Figure S7:** Default values set for the node OptDoE Parts Reference Generator if the user leaves the input empty.

#### 2 Enriched Metabolic Model File: rpSBML

Because some the standard SBML model cannot store strutural information of the chemical species, thermodynamic properties of reactions and pathway information, we use an in-house enriched SBML version that we refer as rpSBML. Using the Minimal Information Required In the Annotation of Models (MIRIAM) conventions, one can store within the SBML file information that instructs the user on the provenance of the reactions and chemical species within the model by cross-references to a wide range of databases. However this needs a third party database lookup to match the database ID with its structural information.

In this work, intermediate products are generated ad-hoc and may not necessarily have database entries. Furthermore, reactions generated by RetroPath2.0 contains information (see additional document) that cannot be contained within the SBML. Similarly, heterologous reactions calculated by RetroPath2.0 contain information on the provenance of the reaction rule, the original reaction used, etc.... That cannot be contained under the MIRIAM annotations as it is. The following information is stored are stored in our enriched SBML version:

- Chemical Species
  - Chemical Structure
    - SMILES
    - InChI
    - InChIkey
- Reactions
  - Reaction rule SMILES
  - Reaction rule ID
  - Reaction rule score
  - Original rule public database ID
  - Path ID

To include such information we elected to enrich the SBML with an in-house annotation labelled "BRSynth". Adding the information to the SBML files in this fashion has the advantage of flexibility to define any new category while maintaining the integrity of the SBML format. Indeed, new annotations enable other parsers to read the file and further analysis may be done with any other software that supports the SBML version 3 format.

#### 3 FBA: Fraction of Reaction

We developed an in-house FBA method to simulate the flux of a target while considering the burden that the production of said target would cause on the cell as a whole. We first perform FBA for the biomass reaction and record its flux. The upper and lower bounds of the biomass reaction are then set to the same amount, defined as a fraction of its previously recorded optimum. This ensures that any further FBA solution would have a fixed biomass production regardless of the conditions set for further analysis. The default fraction is 75% of the optimum, and this value may be changed by the users to define more appropriate burden for the production of a given target molecule would have.

The name of the method is intended to communicate that its more generic than the above description suggests. Indeed, since the user may define the source reaction (or the reaction he would like to restrict) as any reaction within an SBML file, other reactions may be defined to perform the same type of analysis. Lastly, the tool optimises the target molecule and records the flux directly to the SBML file and all changed bounds are reset to their original values before saving the SBML file.

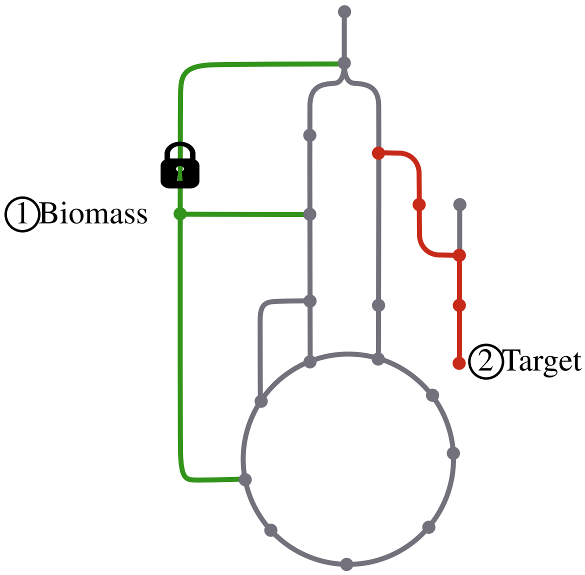

**Figure S8:** Illustration of the "fraction of reaction optimum" method. The whole network depicts a whole cell model, the green lines and circles illustrate the reactions and chemical species that are involved in the biomass reaction and the red are the heterologous ones that produce a target while the grey lines and circles are the original GEM model. The algorithm first optimises the biomass reaction and fixes the bounds of that reaction as 75% of its optimum (illustrated by the black lock). Then the target is optimised with the bounds set for the biomass reaction.

### 4 Pathway Matching Algorithm

This algorithm is designed to quantify the degree of similarity between a measured pathway (from the literature) and a list of predicted pathways using the SynBioCAD platform. Since extracting information from journal articles can be difficult and reports are commonly incomplete, the algorithm reports to what degree of confidence does a measured pathway match with a predicted one. This assumes that the predicted pathway contains the most information while the measured pathway only contains partial information.

Throughout this document, we will refer to $M$ as either the pathway, reaction or chemical species of the measured pathway and $S$ of the predicted pathways.

#### 4.1 Species Match

The first step in the algorithm is to collect and compare the predicted chemical species with the measured ones. To this end a similarity score is computed between the two, by comparing the cross-reference (MIRIAM) annotations of the species and the three layers of their InChI-key in the following manner:

---

**Result:** i
$i \leftarrow 0.4 \cdot \ J(\mathrm{S}_{MIRIAM}, M_{MIRIAM})$
**if** $M_{InChIKey\ Layer\ 1} = S_{InChIKey\ Layer\ 1}$ **then**
    $i \leftarrow i + 0.2$
    **if** $M_{InChIKey\ Layer\ 2} = S_{InChIKey\ Layer\ 2}$ **then**
        $i \leftarrow i + 0.2$
        **if** $M_{InChIKey\ Layer\ 3} = S_{InChIKey\ Layer\ 3}$ **then**
            $i \leftarrow i + 0.2$

---

J($S_{MIRIAM}$, $M_{MIRIAM}$) is the Jaccard index similarity between the MIRIAM annotations, where the MIRIAM annotations are first transformed to matrices before being compared. A perfect match would return a Jaccard score of 1.0 and a completely dissimilar match would return a score of 0.0.

Due to the nature of the species match algorithm, a given measured chemical species can match multiple predicted chemical species. However, we know that a measured chemical species must match only a single predicted one. To make sure that the match is unique, a given measured chemical species is compared with all the chemical species in the predicted pathway and only the best matching one is kept. To illustrate how the algorithm works, consider the case where a measured pathway contains three different chemical species that we compare with a pathway of four, and using the above algorithm, a matching score is computed:

$$\begin{array}{c} \\ S_1 \\ S_2 \\ S_3 \\ S_4 \\ S_5 \end{array} \begin{array}{c} \begin{array}{ccc} M_1 & M_2 & M_3 \end{array} \\ \begin{bmatrix} 0.1 & 0.6 & \boxed{1.0} \\ 0.0 & 0.0 & 0.0 \\ 0.0 & \boxed{0.6} & 0.0 \\ 0.0 & \boxed{0.6} & 0.0 \\ \boxed{0.5} & 0.0 & 0.8 \end{bmatrix} \end{array} \tag{1}$$

For each measured chemical species, the algorithm selects for the best predicted chemical species, preserving the uniqueness of a match. In the above example, the result would be $M_1 \rightarrow S_5$, $M_2 \rightarrow S_3$, $S_4$ and $M_3 \rightarrow S_1$ (highlighted in blue).

Ultimately, the chemical species match is performed to compare two reactions and determine if they are the same by matching all the reactants and products of a reaction with. When a given measured species matched with more than one predicted species with the same score (or more), that information is kept.

#### 4.2 Reaction Match

To compare two reactions $M$ and $S$, all the reactants from a measured reaction ($M_r$) are compared with all reactants from a simulated reaction ($S_r$), and similarly for the products ($M_p$ and $S_p$). Thereafter the reaction score is computed based on all the best scoring chemical species matches:

$$R_s = \overline{\top M_r \cap S_r \; and \; \top M_p \cap S_p} \tag{2}$$

Where the $\top$ represents the top species match for all the intersection $\cap$ between the reactants of products of a reaction. As mentioned in the previous section, when multiple predicted species match with a measured species, the algorithm assumes uniqueness of a match in the scope of the reaction and attempts to filter already matched species. $\top$ represents the top species match for all the intersection $\cap$ between the reactants of products of a reaction. As mentioned in the previous section, when multiple predicted species match with a measured species, the algorithm assumes uniqueness of a match in the scope of the reaction and attempts to filter already matched species. And if that fails, the algorithm assumes that both can be a possibility where all are tested.

Then the EC numbers of the reaction are compared such that:

```
Result: i
i ← 0.0
if M_ec1 = S_ec1 then
    i ← i + 0.25
    if M_ec2 = S_ec2 then
        i ← i + 0.25
        if M_ec3 = S_ec3 then
            i ← i + 0.25
            if M_ec4 = S_ec4 then
                i ← i + 0.25
```

where only the best matching EC number comparison is kept in the score.

Lastly, the two scores are combined with a weighted mean, such that:

$$r = (0.8, 0.2).(R_s, EC)^T \tag{3}$$

Similarly to the species match, a measured reaction can match multiple predicted ones and thus the matches are computed in a matrix and the algorithm selects the best one. Unlike the species matching algorithm, when two reactions have the same score the algorithm assumes still computes a score but flags the reaction as not found.

#### 4.3 Pathway Match

Because predicted pathways can be of different lengths to the measured one, we define a pathway length penalty score:

$$h = 1.0 - \frac{|M_{length} - S_{length}|}{L} \tag{4}$$

Where L is the maximal value that $M_{length}$ and $S_{length}$ can have. If two pathways have the same number of steps $h$ would return 1.0. The penalty is applied to the sum of the reaction match score giving the final pathway match score:

$$f = \overline{r_m} * h \tag{5}$$

where $m$ are all the reactions in a pathway.

#### 4.4 Multiple Pathways Match

When running the SynBioCAD workflow, more than one possible pathway is commonly reported. The result of the pathway matching algorithm is a list (sequence) ordered by $f$:

$$K = f(S_1, M), f(S_2, M), \cdots, f(S_n, M) \quad (6)$$

Where $n$ is the number of pathways predicted by RetroPath2.0. Note that if $f(S_n, M) = 0.0$ or below the threshold (see Figure S9) then that pathway is not included in the list; since its order within the ranked list cannot be ascertained.

##### 4.5 Matching Score Threshold

To define what is the best threshold matching score, we performed the following analysis on 81 measured pathways as reported in the literature (see XLS in additional information). First the best scoring pathway for each collection of predicted pathways with a measured pathway are computed (blue line in Figure S9). A boolean flag is set to inform if all the measured reactions are accounted for within the best scoring predicted pathway (shown in red is Figure S9. This includes predicted pathways that are longer than the measured one. In such a case, a penalty score is imposed on the matching score). Using this, a threshold is determined where any matching pathways with a score lower than that value are ignored. A conservative threshold would be 0.4 where all the matched pathways account for all the measured reactions. However, we use a more loose threshold of 0.25 that lets some of the reactions where the majority of the pathways (47 out of 81 pathways) .

Furthermore, when comparing the literature pathway with the collection of predicted pathways generated by RetroPath2.0, the best similarity scores among them can either be the same or close to best such that more than one of the predicted pathways match with the literature pathway.

#### 5 Global Score

In this section, we look into detail how the global score is computed after pathway analysis in the SynBioCAD platform. This involves normalisation of the different pathway criteria and then performing a weighted mean of the results. The weights are to be optimised based on the comparison between measured and simulated pathways.

##### 5.1 Parameter Normalisation

The following parameters are normalised:

- Pathway Gibbs free energy
- Target flux with fixed biomass flux
- Length of pathway

To enable the comparison between parameters we use the following derivative of the min-max feature scaling with:

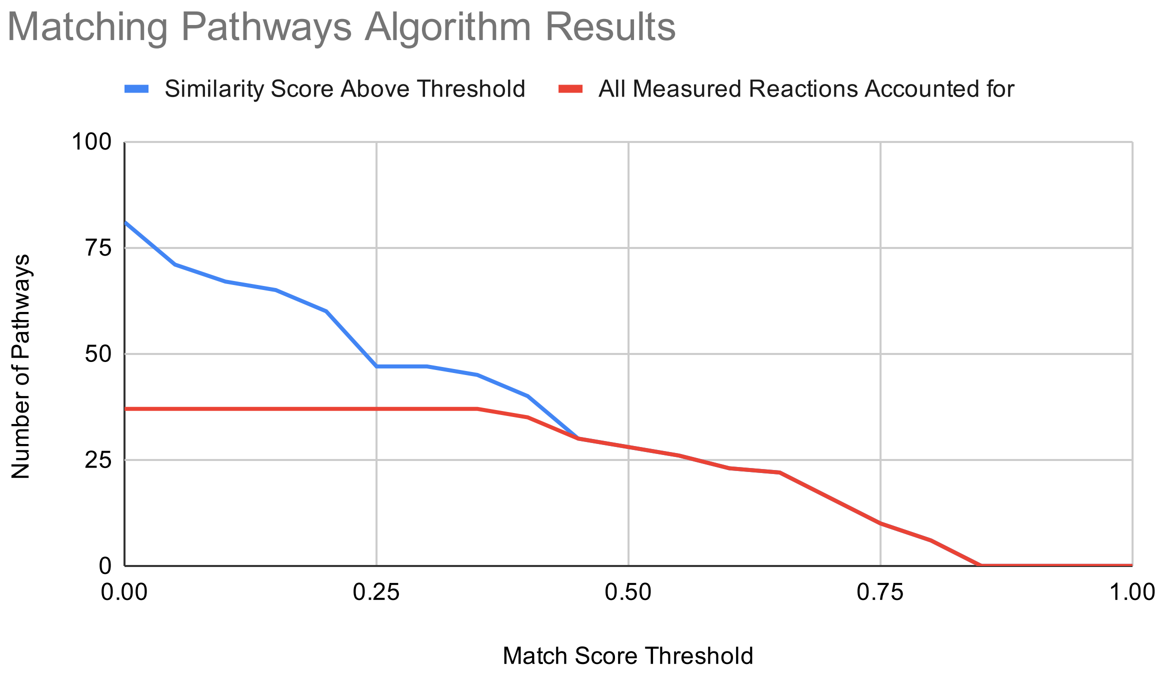

**Figure S9:** Results of the pathway matching algorithm given a range of different matching score thresholds. The blue line represents all the pathways that have a score that is equal or higher to the specified similarity score threshold as shown on the x-axis. The red line shows the number of these pathways that have all of their reactions accounted for. At its best, from a start of 81 literature pathways, the algorithm finds 37 pathways where all the reactions are accounted for by either full species match for all reactions or EC match. The matching threshold used in this study is 0.25 where 47 collection of predicted pathways have at least one pathway that matches with the literature pathway above the threshold.

$$x' = \frac{x - \lfloor x \rfloor}{\lceil x \rceil - \lfloor x \rfloor} \tag{7}$$

Where:

$$x' = \begin{cases} 1.0 & \text{if } x' > 1.0 \\ 0.0 & \text{if } x' < 0.0 \end{cases} \tag{8}$$

##### 5.1.1 Thermodynamics

Because we know the distribution of Gibbs free energy for individual chemical species but not for reactions or pathways, we normalise the species and use the average to determine the pathway thermodynamics.

$$\sum_{n=1}^{n} \Delta_r G_n'^m \tag{9}$$

Where the ceiling and floors are:

$$\begin{aligned} \lfloor \Delta_r G'^m \rfloor &= -7570.2 \\ \lceil \Delta_r G'^m \rceil &= 8901.2 \end{aligned} \tag{10}$$

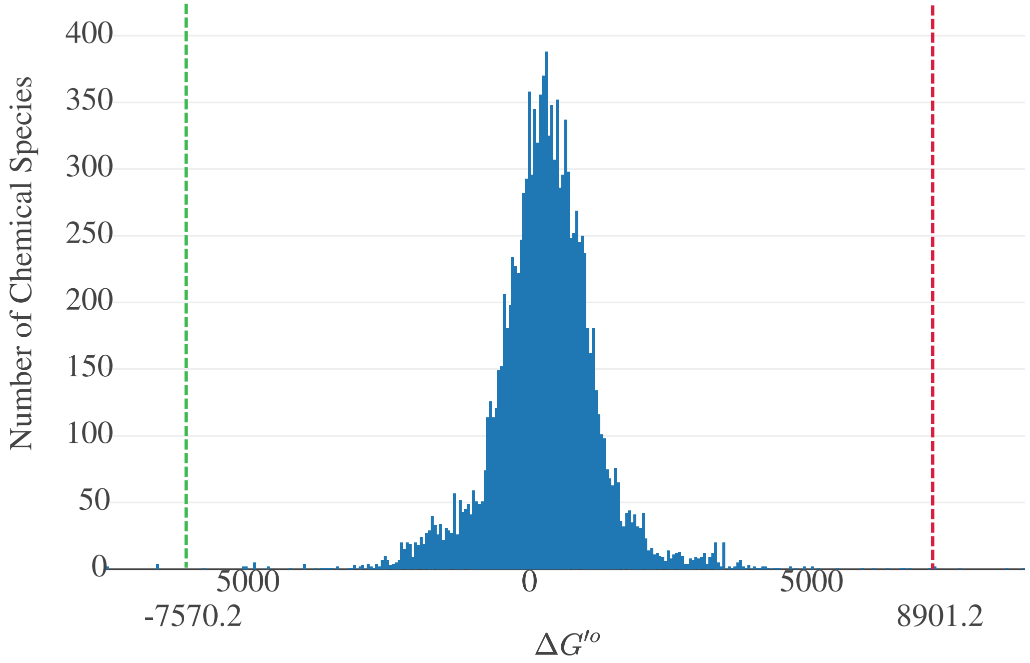

**Figure S10:** All chemical species formation energy contained within the http://equilibrator.weizmann.ac.il/ site. The histogram was used to define the maximal (red) and minimal (green) bounds for normalisation.

##### 5.1.2 FBA

Pathway flux for the target is normalised using the following values, where $j$ is the flux:

$$\begin{aligned} \lfloor j \rfloor &= 0.0 \\ \lceil j \rceil &= 5.0 \end{aligned} \tag{11}$$

###### 5.1.3 Length of Pathway

Since the pathway length is relative to the simulated maximal length, we use the RetroPath2.0 maximal length to define what is the upper limit. We denote the length of the pathway $l$ such that:

$$\begin{aligned} \lfloor l \rfloor &= 0 \\ \lceil l \rceil &= \text{Simulated Maximal Length} \end{aligned} \tag{12}$$

##### 5.2 Weighted Mean

To calculate the global score, the Gibbs free energy, the flux and the length of the pathway are normalised (as defined the in the previous section). We also use mean score from RetroRules ($rr$) over the confidence in the reaction (See RetroPath2.0 original paper) for all the reactions in a pathway as an input to the global score.

We use a weighted mean, where the weights are optimised to literature pathways:

$$(w_j, w_{\Delta G}, w_l, w_{rr}) \cdot (j', \Delta_r G'^{m\prime}, l', rr')^T \tag{13}$$

The weights are to be optimised when comparing measured with simulated pathways and the $'$ denotes the normalised values for each of these criteria.

#### 6 Optimisation

The SynBioCAD platform predicts a number of heterologous pathways to produce a compound of interest. Given a known (measured) pathway, and a list of predicted pathways, here we wish to optimise the weights associated with the following criteria associated with a metabolic pathway: Flux of the target, Gibbs free energy of the pathway, length of the pathway and the reaction rule score. Ultimately we wish to reproduce the order by which the measured pathway matches with the predicted ones.

The literature dataset may be found here.

Given a set of weights, the first step in the optimisation, is to generate an ordered list based on the global score (the global score function is defined as $g$):

$$L(w_j, w_{\Delta G}, w_l, w_{rr}) = \overline{g(S_1, w_j, w_{\Delta G}, w_l, w_{rr}), \cdots, g(S_n, w_j, w_{\Delta G}, w_l, w_{rr})} \tag{14}$$

Where $n$ is the number of pathways returned by RetroPath2.0. The resulting ranked list ($L$) is to be compared with the ranked list generated by the ranked pathway ($K$). Assuming that the measured pathway represents the best pathway, the weights associated with the four different criteria are optimised to have the ranked pathway reflect the best scoring pathways at the top of the rank.

To compare multiple pathways, we use the Rank-Biased Overlap (RBO) between the ranked list of matched pathway (that contains the order we would like to reproduce) and the simulated list of pathways with their global score calculated (where the weights we optimise to match the measured pathway matched ranked list). The main advantage of RBO over other ranked correlation algorithms such as Kendall's $\tau$ or Spearman's $\rho$, is the possibility of comparing two lists of disparate sizes. Indeed, the matching algorithm may return multiple pathways that have a score of 0.0 whose order in the ranked list cannot be ascertained. These such pathways are filtered from the matching ranked list.

$$RBO(K, L(w_j, w_{\Delta G}, w_l, w_{rr}), p) \tag{15}$$

To define $p$, we used the top weightness of the RBO to cover as much as possible before getting the correct one, and was set to $0.9$.

The optimisation problem can thus be posed as the following:

$$\begin{aligned} \text{maximize} \quad & \overline{RBO(K_m, L_m(w_j, w_{\Delta G}, w_l, w_{rr}), p)} \\ \text{subject to} \quad & 1.0 \geq w_j > 0.0, \\ & 1.0 \geq w_{\Delta G} > 0.0, \\ & 1.0 \geq w_l > 0.0, \\ & 1.0 \geq w_{rr} > 0.0 \\ & \sum \{w_j, w_{\Delta G}, w_l, w_{rr}\} = 1.0 \end{aligned}$$

Where $m$ are the number of collection of pathways that are generated by RetroPath2.0 for a given target molecule.

To solve this problem, we used a combination of SOBOL sequencing and genetic algorithm.
